# Coronary artery established through amniote evolution

**DOI:** 10.1101/2022.09.06.506796

**Authors:** Kaoru Mizukami, Hiroki Higashiyama, Yuichiro Arima, Koji Ando, Shigetomo Fukuhara, Sachiko Miyagawa-Tomita, Hiroki Kurihara

## Abstract

Coronary arteries are part of the vascular system that nourishes the heart; they are generally considered a synapomorphy of jawed vertebrates. However, the so-called coronary arteries originated from different body parts in amniotes and other groups, and the evolution of these arteries remains unclear. Here we propose that the amniote coronary arteries were newly obtained, overriding the ancestral arterial systems. In mouse (*Mus musculus*) and quail (*Coturnix japonica*) embryos, the amniote-type coronary arteries are established by the reconstitution of the transient vascular plexus (aortic subepicardial vessels; ASVs) on the outflow tract and the primitive coronary plexus during the development. In contrast, amphibians (*Xenopus laevis, Hyla japonica, Lithobates catesbeianus*, and *Cynops pyrrhogaster*) retain the ASV-like vasculature as extrinsic cardiac arteries throughout their lives and have no primitive coronary plexus. A comparison of zebrafish (*Danio rerio*) and chondrichthyans (*Lamna* sp., *Narke japonica*, and *Deania calcea*) suggested that their hypobranchial arteries correspond morphologically to the ASVs and also serve as heart-feeding arteries throughout their lives. Thus, the coronary artery of adult amniotes is an evolutionary novelty that has acquired new anatomical connections through the addition of a new developmental process to the ancestral pattern. This change is probably related to the modification of branchial arteries, highlights the drastic morphological changes underlying the physiological transition in amniote evolution.

## INTRODUCTION

Coronary arteries are blood vessels that supply oxygen and nutrients to the heart muscle. They are thought to be a synapomorphy of jawed vertebrates because they are absent in the cyclostomes (lampreys and hagfishes) (Grant and Regnier, 1926). In humans, the disturbances in coronary flow, caused mainly by atherosclerosis and associated thrombosis, result in coronary artery disease, the leading cause of death throughout the world (Nabel and Braunwald, 2012; Khera and Kathiresan, 2017). Anatomical similarities in the origin and distribution of the coronary arteries among extant amniotes corroborate the functional importance of these arteries. Among non-amniote groups, coronary arteries are morphologically diverse and are frequently lost in fishes, indicating that they are less important in these animals (see Grant and Regnier, 1926). These differences suggest that coronary arteries are essential for survival in amniotes. However, what morphological changes have occurred in the evolution of amniote coronary arteries is unknown.

In amniotes, the branching point of the coronary arteries is located at the aortic sinuses close to the ventricle. In teleosts and chondrichthyans, it is located in more rostral vessels, mainly in the hypobranchial arteries, running long distances toward the heart (Parker, 1886; Grant and Regnier, 1926; Corrington, 1930; May and Herber, 1957; Halpern and May, 1958; Muñoz-Chápuli et al., 1994; Icardo, 2017). Such positional differences have led to two hypotheses: coronary arteries gradually shifted from the dorsal to the ventral position in association with the remodeling of branchial arches during the water-to-land transition (Halpern and May, 1958, MacKinnon and Heatwole, 1981), or non-homologous arteries nourishing the heart are collectively termed “coronary arteries” (Grant and Regnier, 1926; May and Herber, 1957). Data on amphibians may help to address this controversy, although few comparative studies have examined their extrinsic blood vessels, and some authors have mentioned that amphibians lack coronary arteries (Kapuria et al., 2018; Lupu et al., 2020).

In the present study, we compared the anatomy and development of the coronary vasculature to clarify its evolutionary history in various vertebrate lineages.

## RESULTS

### Artery distribution in the heart differs between amniotes and amphibians

We first compared the anatomy in mice (*Mus musculus*), quails (*Coturnix japonica*), and some amphibians. In the 17.5 dpc mouse fetus (*n* = 5), the two coronary arteries branched from the root of the aorta, which is part of the outflow tract (Figs. 1A, S1). They were distributed on the surface of the ventricle. Similarly, in quail at stage 28 (*n* = 4), the two coronary arteries branched from the aortic root and were distributed on the ventricular wall (Fig. 1A). In the adult *Xenopus laevis* (*n* = 6), no blood vessels were found on the ventricular surface. The extrinsic artery was found only on the surface of the outflow tract, whose branching point (orifice) was located in the left aortic trunk, the bifurcated part of the outflow tract (Fig. 1B). Identical topographical relationships were also found in the other frog (*Lithobates catesbeianus*) and newt (*Cynops pyrrhogaster*) (*n* = 3 each; (Fig. 1B). These results indicate that the morphology of so-called coronary arteries is highly divergent between amniotes and amphibians, and it is difficult to compare them in terms of their distribution areas and orifice positions.

**Fig. 1:**
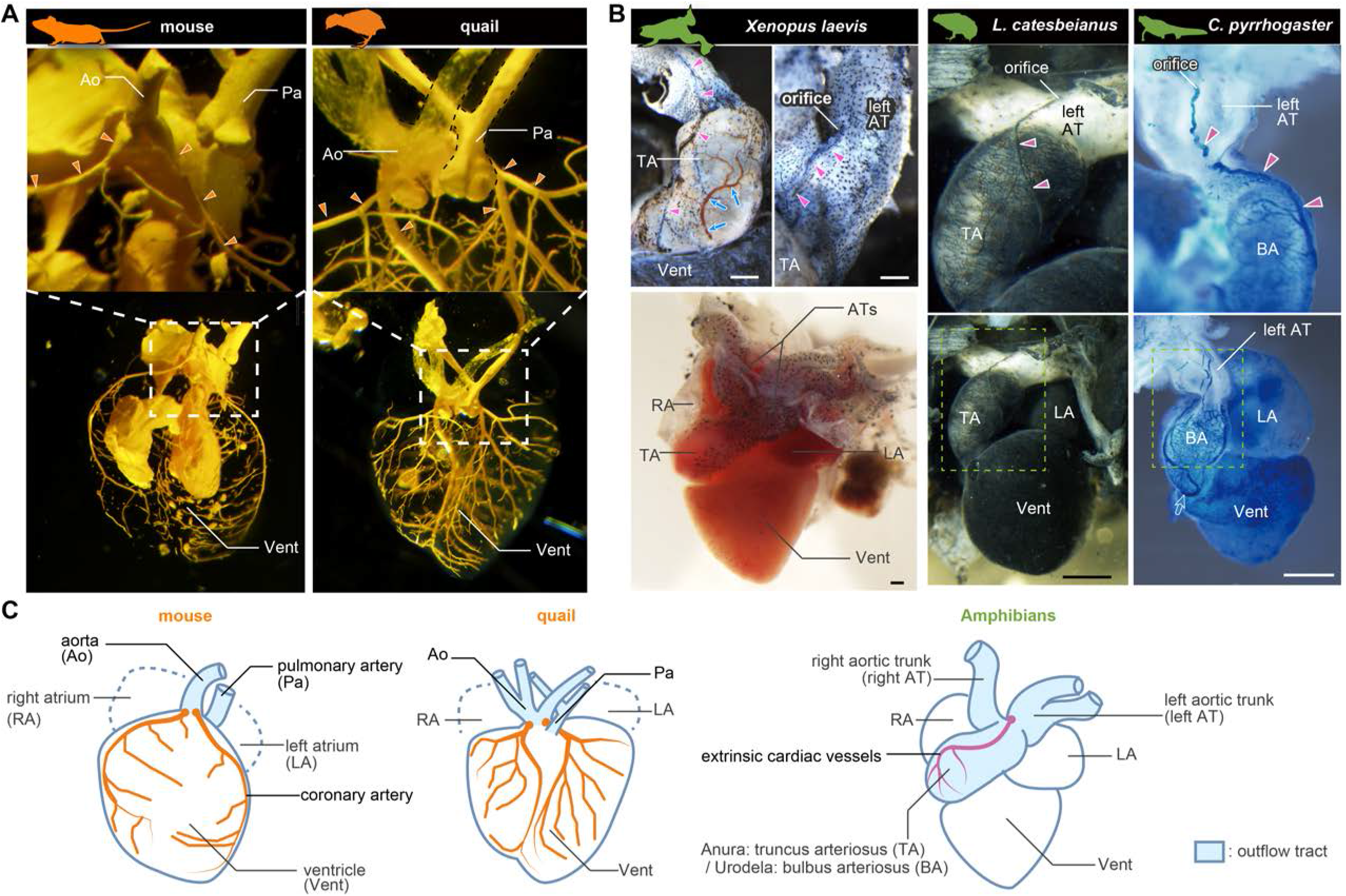
Anatomy of so-called coronary arteries in mouse, quail, and amphibians. **(A)** Resin-injected coronary circulations of mouse (17.5 dpc) and quail (stage 28). **(B)** Ink-injected hearts of *Xenopus* (adult frog), African bullfrog (*Lithobates catesbeianus*; tadpole), and Japanese fire-belly newt (*Cynops pyrrhogaster*; adult). Arrowheads indicate the extrinsic arteries on the outflow tract. No vessels were found in the ventricular wall. **(C)** Summary. Scale bars: 1 mm.

### Coronary arteries develops in two steps in amniotes and in one step in amphibians

In the 11.5 dpc mouse embryo, a primitive coronary plexus was found on the ventricular surface. PECAM-1-positive arteries were observed in the cranial region of the outflow tract, and were located on the ventral side of the third to fifth pharyngeal arches (*n* = 3; Fig. 2A). At 12.5 dpc, the arteries extended from the cranial toward the aortic root with ramification (*n* = 4; Fig. 2A). At 13.5 dpc, the arterial network covered the aortic root and was connected with the primitive coronary plexus. Simultaneously, the cranial part of this vascular trunk was lost (*n* = 7; Fig. 2A). By 15.5 dpc, the true coronary artery was established with a complete loss of the arterial vasculatures on the outflow tract and modifications of the primitive coronary plexus (*n* = 3; Fig. S1).

**Fig. 2:**
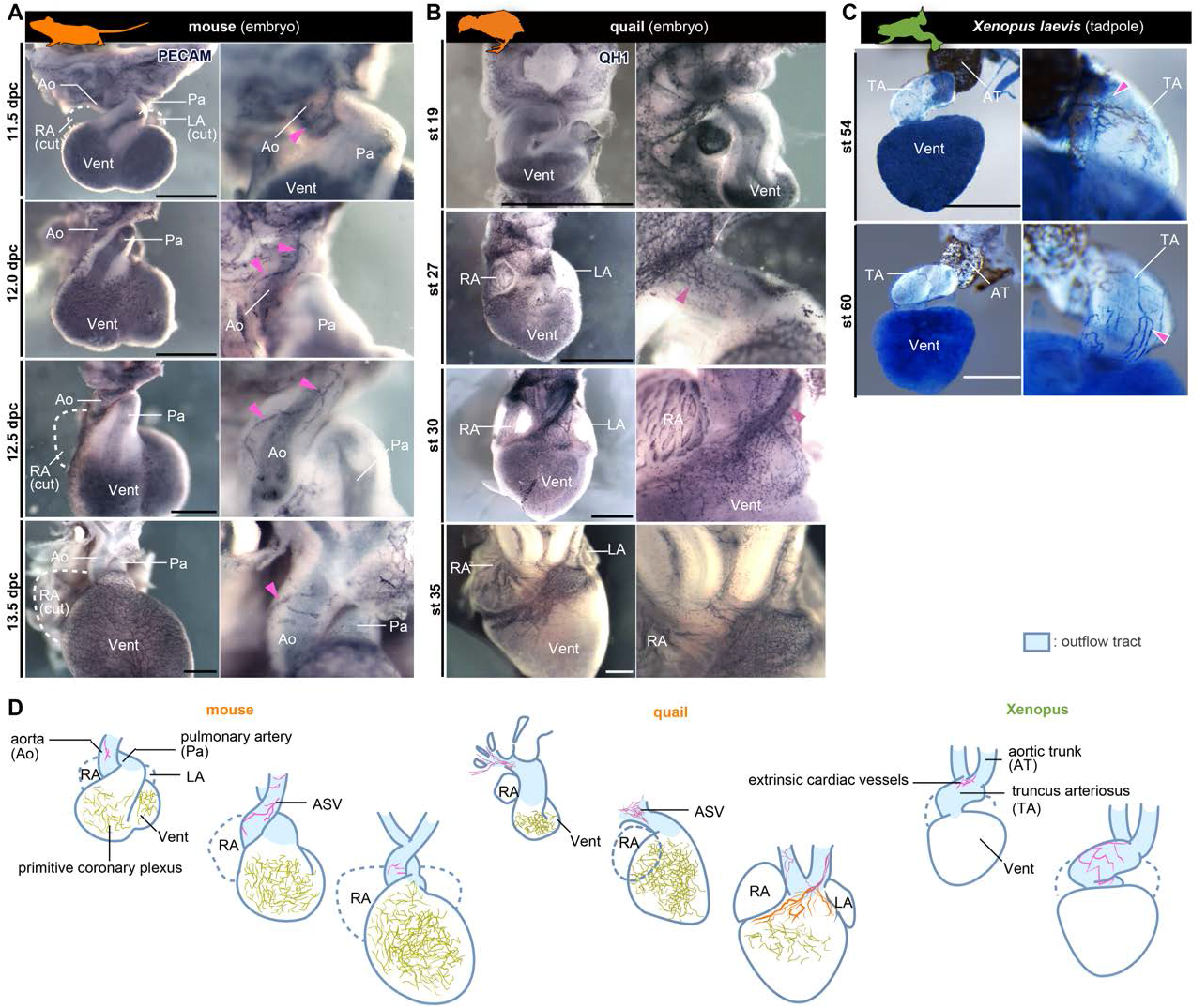
Development of the so-called coronary arteries in mouse and quail embryos and in *Xenopus* tadpole. **(A, B)** Two types of primitive blood vessels, aortic subepicardial vessels (ASVs) and the primitive coronary plexuses were present in the embryonic stages in mouse (A) and quail (B). The blood vessels were visualized by the immunohistochemistry. The pink arrowheads indicate ASVs or the small vessels originated from the cranial part of the outflow tracts. **(C)** Extrinsic cardiac arteries of *Xenopus*. The blood vessels were visualized by the ink injection. **(D)** Summary. LA, left atrium; RA, right atrium. Scale bars: 500 μm.

The development of quail was consistent with that of mouse (Fig. 2B). At stage 19 heart, no QH-1-positive cells were observed on the surface of the outflow tract (*n* = 2) (also see Ainsworth et al., 2010). At stage 27, QH-1-positive endothelial cells formed a fine vascular plexus from the cranial part of the outflow tract toward the aortic root (*n* = 3). At stage 30, the plexus extended around the aorta and pulmonary trunk to form the arterial network (*n* = 4). At stage 35, the arterial plexuses of the aorta and pulmonary trunk degenerated, and the coronary arteries were formed secondarily on the ventricle surface (*n* = 2).

In *Xenopus*, the extrinsic vessels were found in the tadpole stage. At stage 54, the extrinsic cardiac arteries appeared in the cranial part of the outflow tract, ventral to the branchial arches (*n* = 2; Fig. 2C). At stage 60, the arterial plexus expanded on the outflow tract to reach the aortic root. This vascular pattern was maintained throughout the development into the adult stage.

The amniote-type coronary arteries developed in two steps. According to Chen et al. (2014), we will henceforth refer to the vascular plexus on the aorta as the aortic subepicardial vessels (ASVs) (Fig. 2D).

### Amphibian extrinsic cardiac arteries as homologs of amniote ASVs

To understand the homology of the extrinsic cardiac vessels, we tried to identify the exact location of their orifices. In 12.5 dpc mouse embryos, the ASV orifice was found in the carotid artery (Fig. 3A), and contained PECAM-1-positive endothelial cells (*n* = 14; Fig. 3B). This PECAM-1-positive ASV ran through the outflow tract, then entered the aortic root before reaching the ventricle to form a secondary orifice (Fig. 3C). At 13.5 dpc, the orifice in the carotid artery was lost, and the secondary orifice became larger (Fig. 3D). In quail embryos, the endothelial invasion into the aortic wall from the ASV network was observed in the aortic root at stage 30 (*n* = 10; Fig. 3E). These results indicate that the amniote ASVs are formed as a continuous endothelial structure from the pharyngeal region to the vascular network around the aortic root and contribute to creating the secondary orifice of the true coronary artery.

**Fig. 3:**
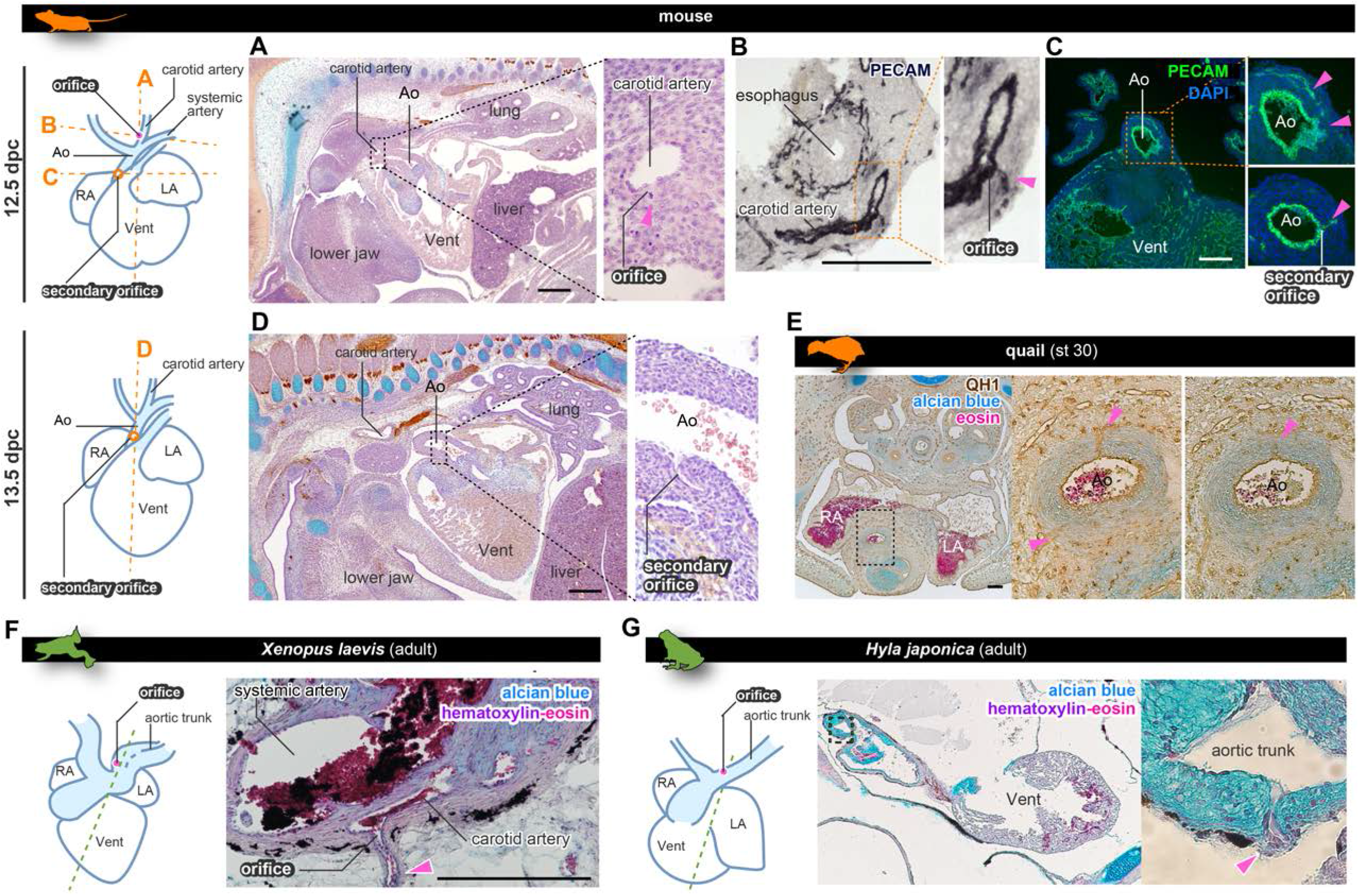
The positions of the orifices of the so-called coronary arteries. **(A–C)** ASVs in 12.5 dpc mouse embryos branched from the future carotid artery. Panel A shows HE staining of paraffin sections, and panels B and C show immunohistochemical staining images using the cryosections. They ran through the outflow tract and formed the secondary orifice in the aortic root close to the ventricle **€**. **(D)** In 13.5 dpc embryos, the secondary orifice developed in the aortic root instead of the carotid artery orifice, which was lost. **(E)** In quails at stage 30, the secondary orifice was formed at the aortic root, as in mice. The immunohistochemical- and HE staining images using the cryosections. **(F)** In adult *Xenopus*, the extrinsic cardiac artery (arrowhead) branched from the carotid artery in the aortic trunk. HE stained paraffin section. **(G)**In the Japanese tree frog (*Hyla japonica*), the extrinsic cardiac artery branched from the aortic trunk close to the root of the carotid artery. HE stained paraffin section. Ao, aorta; LA, left atrium; RA, right atrium; Vent, ventricle. Scale bars: 100 μm **(A–E)**, 1 mm **(F, G)**.

In the adult *Xenopus*, the aortic trunk contained the systemic and carotid arteries; the orifice of the extrinsic cardiac artery opened into the carotid artery (*n* = 2; Fig. 3F). An identical orifice in the Japanese tree frog (*Hyla japonica*) was positioned at the root of the left carotid artery, and vessels extended to the surface of the aortic trunk (*n* = 4; Fig. 3G). We concluded that, because of their topographical relationships, the amphibian extrinsic cardiac arteries are comparable to amniote ASVs rather than to the coronary arteries.

### The coronary vein is conserved among tetrapods

In amniotes, blood supplied by the coronary arteries enters the coronary veins and is collected in one trunk, which opens into the sinus venosus (Grant & Regnier, 1926; Fig. 4A). In 15.5 dpc mouse embryos, the connections of the coronary artery and the vein were established (Fig. S1). Using several amphibian species, we checked whether this vein morphology is conserved in amphibians.

**Fig. 4:**
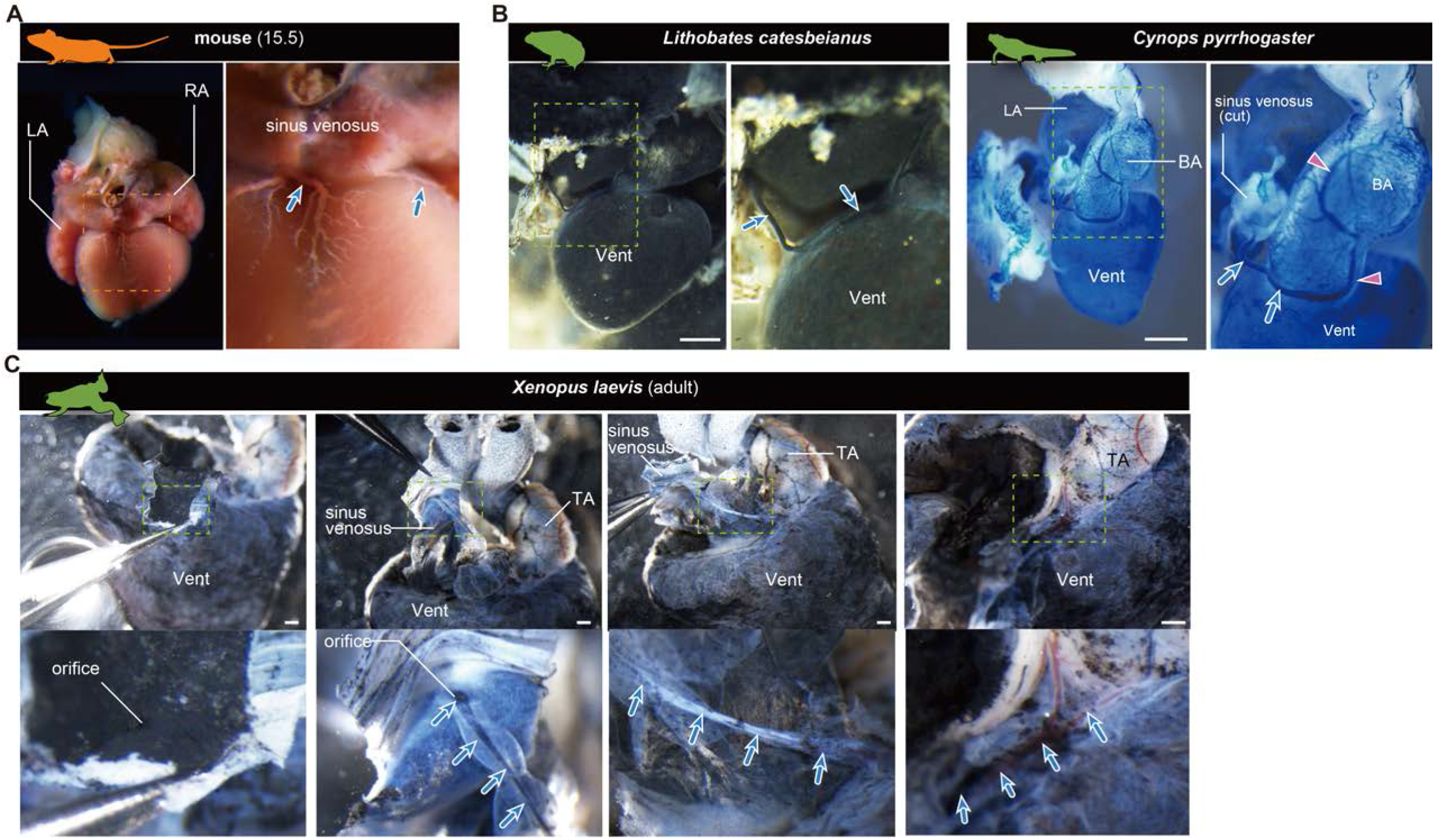
The orifices of the coronary veins are conserved among tetrapods. Ink-injected hearts are shown. **(A)** Mouse. **(B)** American bullfrog(*Lithobates catesbeianus*) and the Japanese fire-belly newt (*Cynops pyrrhogaster*). **(C)** *Xenopus*. Arteries (pink arrowheads) and veins (blue arrows) were found on the surface of the outflow tract. In **(B)** and **(C)**, coronary vein (blue arrows) branched from the sinus venosus and passed to reach the outflow tract. BA, bulbus arteriosus; LA, left atrium; RA, right atrium; TA, truncus arteriosus; Vent, ventricle Scale bars: 1 mm.

In amphibians, the venous drainage from the wall of the outflow tract converged into a single coronary vein, which ran down laterally along the right coronary sulcus (Figs. 1B, 4B, 4C) and opened into the junction between the sinus venosus and the atrium (Fig. 4B, C). The coronary vein was also identified as a connection between the vasculature on the outflow tract and the junction between the sinus venosus and atrium through the right coronary sulcus (Fig. 4C).

Thus, coronary arteries and veins are not distributed on the ventricle in frogs and newts. In amphibians, the ventricle walls are spongy and thin (Grant and Regnier, 1926), so there may be no need to distribute fine blood vessels to supply blood outside the heart. Nevertheless, extraneous arteries and veins are distributed and supply blood, at least in the outflow tract. Although the peripheral distribution is very different from that in amniotes, amphibians have a vascular system equivalent to coronary veins.

### So-called coronary arteries in fishes

To investigate the evolutionary history of coronary arteries, we conducted outgroup comparisons using zebrafish (*Danio rerio*) and chondrichthyans. In adult *flk1:egfp* zebrafish (72 dpf), a peripheral network of coronary vessels was found throughout the entire surface of the ventricle (*n* = 6; Fig. 5A). These “coronary arteries” led to the heart over a long distance directly from the hypobranchial artery, which branched off from the branchial arches (Fig. 5A, B; see also Hu et al., 2001). Before reaching the ventricle, this artery bifurcated, and its two branches extended ventrally and dorsally to connect to the vascular network on the ventricular wall.

**Fig. 5:**
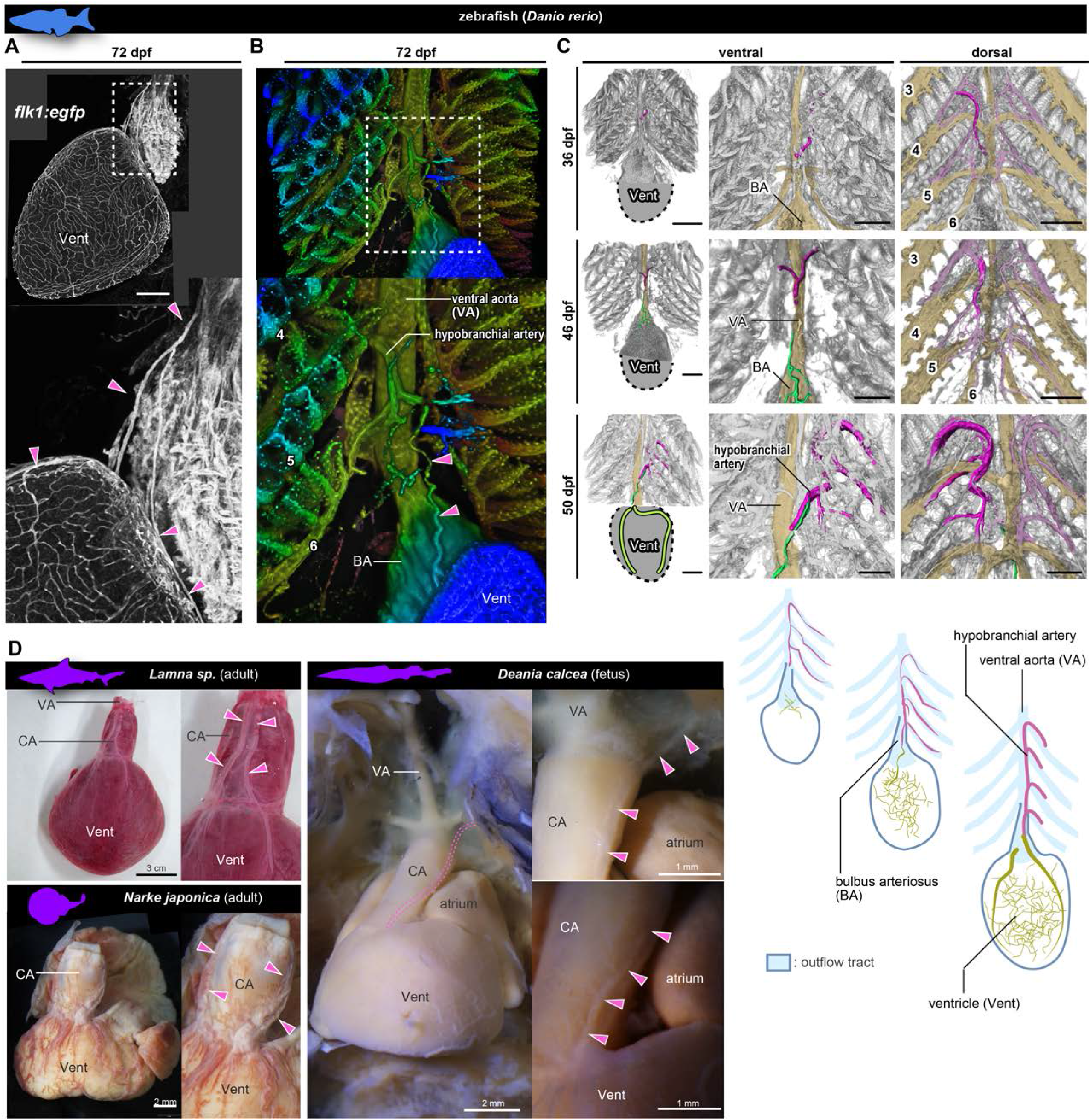
Anatomy and development of the so-called coronary arteries of fishes. **(A, B)** *flk1:egfp* zebrafish at 72 dpf (juvenile). **(B)**Three-dimensional image. (**C**) *flk1:egfp* zebrafish at 36, 46, and 50 dpf. The hypobranchial arteries (pink) arose from the dorsal side of the pharyngeal arch arteries. The vascular network (green) appeared on the ventricular surface at 46 dpf and connected with the hypobranchial artery at 50 dpf. The numbers indicate the branchial arteries. **(D)** Chondrichthyans. The anatomical pattern of the hypobranchial and coronary arteries was identical to that of zebrafish. BA, bubus arteriosus; CA, conus arteriosus; VA, ventral aorta; Vent, ventricle. Scale bars: 100 μm.

At the late pharyngula stage (31–38 hpf) of *flk1:egfp* zebrafish embryos, the Y-shaped hypobranchial artery appeared in the midline of the pharyngeal region, and its lateral branches bilaterally connected with the first pharyngeal arteries of the mandibular arch (*n* = 2; Fig. S2A). The mid-caudal branch of the Y-shaped artery extended caudally toward the cardiac outflow tract. After hatching (48–72 hpf), it further extended along the ventral aorta (3–14 dpf) and formed a vascular plexus on the aorta around 21 dpf (*n* = 10; (Fig. S2A). Around 36 dpf (juvenile stage), the hypobranchial artery was identified as a vessel running on the ventral side of the ventral aorta, but no vascular network was found on the surface of the ventricular wall (*n* = 3; Fig. 5B). At 45 dpf, a vascular plexus was formed on the ventricle close to the aortic root, and part of it covered the outflow tract (Fig. S2B). At 46 dpf, it formed connections with the mid-caudal elongation of the hypobranchial artery (*n* = 13; Figs. 4C, S2B). The “coronary arteries” on the ventricular wall were completely formed by around 50 dpf, slightly before the adult stage.

The relationship between zebrafish hypobranchial artery and the ventricular vascular plexus is similar to that between ASVs and the primitive coronary plexus in amniotes. However, zebrafish does not form a novel orifice in the heart proximity or lose or reconstitute the above vessels. The hypobranchial artery is not a transient structure and is maintained as the trunk of the coronary artery even in adults.

The anatomy of zebrafish-type coronary circulation was also conserved in the Chondrichthyes (Fig. 5D). In *Lamna* sp. (*n* = 1) and *Narke japonica* (*n* = 3), the coronary artery originated from a vessel further cephalad than the bulbus arteriosus (outflow tract). In *Deania calcea* (*n* = 3), the arterial trunk branched from the branchial artery, bifurcated on the conus arteriosus (outflow tract), and then supplied its peripheral branches to the ventricle.

## DISCUSSION

### ASV as the ancestral component of the coronary artery

Since Ibn al-Nafis discovered coronary arteries in the 13^th^ century (Numan, 2014), their developmental origin has been an enigma. In amniotes, coronary arteries had been traditionally thought to originate from the aortic wall by outgrowth, sprouting angiogenesis (Bennett, 1936; Hutchins et al., 1988). However, we found that the amniote-type coronary arteries developed by the reconstitution of primary vessels. This is consistent with the previous studies showing the ingrowth formation of amniote orifices in the connections between ASVs and primitive coronary plexus (Bogers et al., 1989; Red-Horse et al., 2010; Peng et al., 2013; Tian et al., 2013; Chen et al., 2014; Ivins et al., 2015; He and Zhou, 2018).

Our data suggest that the amniote ASVs originated from the pharyngeal region (particularly the arches posterior to the third pharyngeal arch) and extended to the aortic root to form the vascular network. This is reminiscent of the contribution of the cardiopharyngeal mesoderm to the heart development (Diogo et al., 2015). Our suggestion that ASVs are derived from the pharyngeal arch is consistent with the data obtained by Maruyama et al. (2019), who found that the periaortic lymphatic endothelium overlapping with ASVs originated from the *Islet1* -positive pharyngeal core mesoderm. Still, we cannot tell the extent to which ASV distribution correlates with the contribution of the cardiopharyngeal mesoderm to the heart. If the neural crest cells and mesoderm of the pharyngeal arches are maintained as developmental modules even in the heart, the ASV distribution may reflect the mesenchyme distribution, just as the cranial nerve distribution follows the distribution of neural crest cells from the facial primordia (Higashiyama et al., 2021).

The so-called coronary artery of amphibians has been described ambiguously as noted in the introduction; we found that amphibian extrinsic cardiac arteries are a system composed solely of ASVs that do not give rise to the primitive coronary plexus on the ventricle. In fishes, it seems reasonable to consider the hypobranchial arteries as vessels comparable to ASVs, since these arteries have an orifice in the pharyngeal arch artery and are distributed along the ventral aorta to the aortic root (Fig. 5C). In zebrafish, Paffett-Lugassy et al. (2017) reported that the hypobranchial artery endothelium is derived from the second heart field in the pharyngeal arch, indicating that both ASVs and hypobranchial arteries share the same cell population from the same body part. Thus, here we propose that the hypobranchial artery of fishes is the ancestral homolog of ASVs in tetrapods.

The above considerations suggest that ASVs were established as hypobranchial arteries in ancestral jawed vertebrates. With branchial evolution and changes in the primitive coronary plexus, this highly conserved component has led to the derivation of various forms of coronary arteries in vertebrate history. Thus, the amniote-type true coronary arteries were formed by overriding the embryonic or ancestral pattern of cardiac arteries.

### Diversity of coronary arteries among vertebrates

The morphology of the coronary arteries of amniotes is relatively uniform, whereas that of non-amniotes is highly diverse and depends on the lifestyle of each species (see Grant and Regnier, 1926). For example, caecilians possess a unique coronary circulation, which is unlike that in any other vertebrate group. In *Hypogeophis rostratus*, the so-called coronary arteries supply the heart tissue arising from the trabecular spaces of the ventricle (Lawson, 1966), while *Epicrionops* reportedly has no coronary arteries (Wilkinson, 1996). Despite this diversity of coronary arteries, the coronary veins of caecilians branch from the sinus venosus (Lawson, 1966; Wilkinson, 1996), as observed in mice, frogs, and newts (Fig. 4).

The coronary vasculature is most variable in teleosts. Fish hearts are often classified into four types (usually termed Type I to IV) on the basis of histological distribution of coronary arteries (Tota et al., 1983; Tota, 1989; Icardo, 2017). Many teleosts even lack coronary circulation, but this must be a derived condition because non-teleost actinopterygians and chondrichthyans have coronary arteries (Farrell et al., 2012; Farrell and Smith, 2017). The lack of coronary circulation is probably caused by the lower need in external oxygenation due to the acquisition of spongy heart walls. Even in chondrichthyans, coronary arteries do not always have ancestral morphology; the “posterior coronary arteries” develop as branches of the subclavian arteries in some stingrays (Muñoz-Chápuli et al., 1994).

Because non-amniote coronary arteries are highly variable among species, it is difficult to draw a fixed anatomical scheme. In amniotes, they are not as diverse (except in turtles; see below), probably because their malformation is associated with lethality. Thus, in ancestral jawed vertebrates, the ASV (hypobranchial artery) and primitive coronary plexus were associated in various ways to give rise to diverse cardiac arteries, but in amniotes, they develop uniformly constrained way with the addition of new developmental processes which remodeling of the ASVs and primitive coronary plexus.

However, the origin of coronary endothelial cells varies among taxa, even though the coronary endothelium is the same histologically (Table 1 in Kapuria et al., 2018). In zebrafish, ventricular circulation is provided by the endocardium of the atrioventricular junction (Harrison et al., 2015). In gourami (*Trichogaster trichopterus*), it is derived solely from the hypobranchial arteries (Shifatu et al., 2018). In the giant danio (*Devario malabaricus*), it arises from the hypobranchial artery and the atrioventricular junction (Shifatu et al., 2018). The developmental origins of the endothelial cells of adult coronary vasculatures vary not only among teleosts, but also among vertebrates and even among amniotes (see Kapuria et al., 2018), indicating that the relative contributions of ASVs and the primitive coronary plexus to adult coronary arteries varies among jawed vertebrates. Or, perhaps, the cellular origin of the primitive coronary plexus is not uniform; the only common feature appears to be undifferentiated vascular vessels distributed in the embryonic ventricle. The differences in cell lineage should not affect artery identity, and it is appropriate to regard them as a developmental system drift, as in the case of developmental modifications in the formation of the nematode vulva (Wang and Sommer, 2011). Anyway, the origin of the ancestral coronary arteries of jawed vertebrates can be defined only by the presence of ASVs and the primitive coronary plexus, and it is not possible to uniquely depict which cells they originated from and how these cells were related to each other to give rise to their coronary vasculatures. In amniotes, a novel developmental process—secondary orifice formation—was acquired from these diverse ancestral developmental patterns, establishing the true coronary arteries essential for survival. Since it can be defined morphologically as a structure with novel connections and is functionally important enough so that its anomaly is immediately lethal, the amniote-type coronary artery should be considered an evolutionary novelty, deviating from the ancestral coronary vasculature.

### Amniote evolution and the establishment of the true coronary artery

The evolutionary history of coronary arteries is summarized in Figure 6A. The acquisition of the amniote-type coronary artery should be related to a drastic reorganization of pharyngeal structures or physiological changes during amniote evolution.

**Fig. 6:**
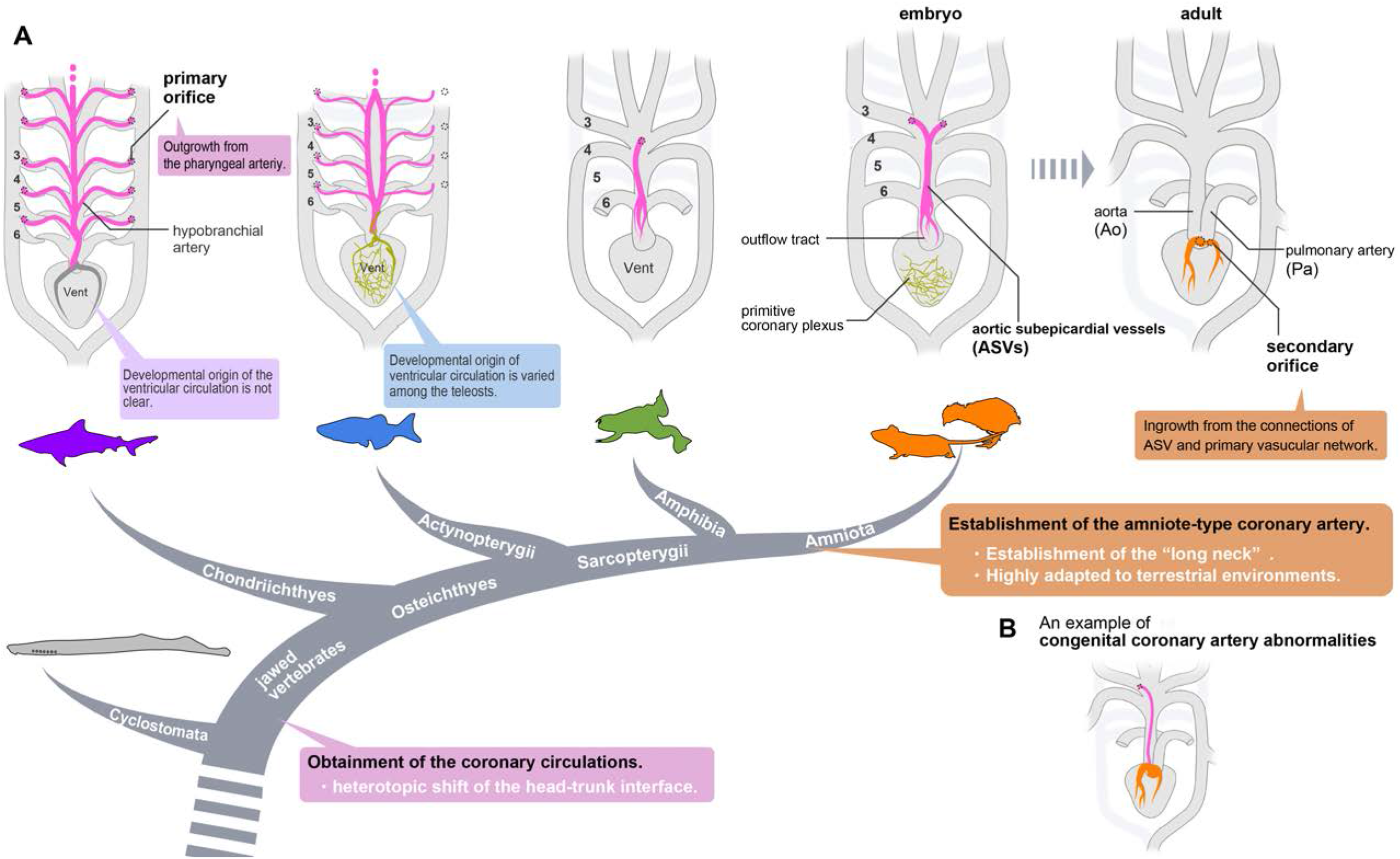
Evolution of extrinsic cardiac arteries. **(A)** The amniote-type coronary arteries are reconstituted from the ASVs and the primitive coronary plexus. The extrinsic cardiac arteries of amphibians and the hypobranchial arteries of fishes are comparable to the ASVs, rather than to the amniote-type coronary arteries. It remains uncertain whether the ventricular artery in chondrichthyans arises as an extension of the hypobranchial artery or has a different developmental origin. As ventricular vessels have a variety of developmental sources in teleosts, ancestrally, the vessels in this region may have had a variety of developmental origins in different animal species. In amniotes, such redundant coronary arterial development gave way to a single robust and unchangeable developmental program, resulting in developmental constraints. **(B)** A scheme of congenital coronary artery abnormalities according to Kim et al. (2009).

The cardiopharyngeal structures in tetrapods have progressively changed during the water-to-land transition to form the amniote bauplan adapted to arid land. Branchial arches have changed into neck structures, and a long neck has formed in the amniotes with the morpho-functional separation of the head and pectoral girdle (Hirasawa et al., 2016). Simultaneously, the heart, located ventral to the pharynx, moved back into the thorax and became separated from the pharyngeal structures (Hirasawa et al., 2016). This heterotopic shift of the heart inevitably resulted in various reorganizations of the pharyngeal arch derivatives, such as the recurrent laryngeal nerves of the vagus, which have been modified into the peculiar route without altering their anatomical connections (Higashiyama et al., 2016a). The establishment of true coronary arteries with a secondary orifice was probably inevitable to achieve both morpho-functional reduction of the branchial structures and the development of a thickened myocardial layer during this cardiopharyngeal modification.

Whether the reorganization of coronary arteries was physiologically essential or not remains unclear. Coronary arteries in lizards and snakes are identical to those in mammals and birds, but the orifice in turtles is in the middle of the carotid artery, as it is in frogs (Grant and Regnier, 1926), suggests that the amniote-type coronary arteries were lost secondarily in the turtle lineage. Thus, transient ASV formation is conserved during development even after acquisition of true coronary arteries, enabling them to return from the amniote type to the ancestral type. This might possibly be related to the peculiar morphogenesis of the shoulder girdle in turtles (Nagashima et al., 2009). The extent to which this structural change has physiological consequences that extend to cardiac function remains unknown.

The evolutionary origin of extrinsic cardiac arteries is still a mystery. Cyclostomes do not have coronary circulation, but whether this is the ancestral condition or a secondary loss is uncertain (Grant and Regnier, 1926). If they have retained the ancestral pattern, the establishment of extrinsic cardiac vessels may be related to the heterotopic shift of the cardiac region and head–trunk interface in the ancestor of gnathostomes (Higashiyama et al., 2016a), although further studies are warranted in this lineage.

### Clinical implications of the coronary development

Using the present theory, we can explain some congenital coronary artery abnormalities, an uncommon disease entity covering a broad spectrum of abnormalities (Fig. 6B). Although many cases remain asymptomatic and are incidentally found by coronary angiography, these abnormalities sometimes cause sudden death (Taylor et al., 1992; Yuan, 2014; Villa et al., 2016). In particular, the anomalous origin of the coronary artery from the innominate (brachiocephalic) artery or adjacent aortic arch is often lethal, accompanied by other severe anomalies (Davis and Lie, 1977; Asada et al., 2019; Pandey et al., 2019), although some patients survive (Santucci et al., 2001; Duran et al., 2008; Kim et al., 2009). Since the brachiocephalic artery is derived from the right third pharyngeal arch artery and they pass across the ventral side of the ascending aorta, the anomalous coronary arteries in these reports are similar to ASVs. Thus, this type of anomaly can be considered as persistent embryonic vasculature.

In conclusion, the present study shows that the true (amniote-type) coronary artery first developed by reconstitution of the ancestral cardiac vessels in amniote evolution; these arteries characterize the amniote heart morphologically and functionally. Our reconstruction of the evolutionary history and morphological redefinition of coronary circulation will be helpful for the evolutionary studies of amniotes and comparisons among animal models in coronary artery disease studies.

## MATERIALS AND METHODS

### Animals

Wild-type mice (*Mus musculus*) of the ICR background were kept in an environmentally controlled room at 23±2°C with a relative humidity of 50–60% and under a 12 h light: 12 h dark cycle. Embryonic ages were determined by timed mating, with the day of the plug being 0.5 days post-coitum (dpc).

Eggs of Japanese quails (*Coturnix japonica*) were obtained from Motoki Hatchery (Saitama, Japan). They were incubated in a humidified atmosphere at 37°C until the embryos reached appropriate stages (Ainsworth et al., 2010).

African clawed frogs (*Xenopus laevis*) were obtained from the Watanabe *Xenopus*-inbred strain resource center (Hyogo, Japan) and were kept at room temperature (RT). The embryonic stage was determined as in Nieuwkoop et al. (1994). Japanese tree frogs (*Hyla japonica*) were caught in Tsukuba (Ibaraki, Japan), and American bullfrogs (*Lithobates catesbeianus*) were caught in Oyamadairi park (Hachioji, Tokyo, Japan) according to the laws of the Invasive Alien Species Act. For Japanese fire-belly newts (*Cynops pyrrhogaster*), we purchased in store Kingyozaka (Tokyo, Japan) to use artificially bred and commercially distributed newts.

Zebrafish (*Danio rerio*) were maintained as described by Fukuhara et al. (2014). Other than the wild-type zebrafish, we used *flk1:egfp* [Tg(*kdrl:egfp*)^s843^; ZFIN: ZDB-ALT-050916-14; Jin et al., (2015)]. Embryos and larvae were staged by hour post-fertilization (hpf) and days post-fertilization (dpf) at 28–28.5°C (Kimmel et al., 1995).

Chondrichthyan samples were obtained through fish markets, mainly from incidental bycatch. The hearts of salmon shark (*Lamna* sp.) were obtained from Yoshiike (Tokyo, Japan), Japanese sleeper rays (*Narke japonica*) were from Induka Shouten (Nagasaki, Japan), and the fetuses of bird-beak dogfish (*Deania calcea*) were from Gyosai-ya (Tokyo, Japan).

The sample size was designed to minimize animal sacrifice as much as possible. We basically considered n = 3 to be sufficient for each developmental stage. Still, we observed even larger numbers of animals, such as mice, for which many embryos were available from a single female. In some experiments, we found n = 2 or less to be sufficient. Except for apparent failures in the experiment (e.g., no signal, excessive background), all other cases were considered as the target samples.

All animal experiments were approved by the University of Tokyo Animal Care and Use Committee (approval ID: P19-043 and P19-050) and were performed in accordance with the institutional guidelines and the Act on Welfare and Management of Animals. We also follow the ARRIVE guidelines for animal research (Percie du Sert et al., 2020).

### Histological sections

For the cryosections, samples were fixed in 4% paraformaldehyde in PBS (4% PFA/PBS) overnight at 4°C and washed with PBS. The samples were transferred into 10% sucrose/PBS, 30% sucrose/PBS, and 30% sucrose in Optimal Cutting Temperature (OCT) compound (Tissue Tek, Sakura Finetek, Japan) for 30 minutes at 4°C for each step, and embedded in 100% OCT compound. The embedded block was stored at −20°C before use. The samples were cryo-sectioned (8-12 μm thick). For the paraffin sections, samples were fixed with Serra’s solution (A mixture of formalin, ethanol, and acetic acid at 7:2:1) overnight and stored in the 70% ethanol at RT. The samples were dehydrated stepwise with ethanol to xylene series and embedded in paraffin. The paraffine blocks were sections in 6-9μm thick. The paraffine sections were stained by alcian blue, hematoxilin, and eosin by using the standard protocol (HE staining).

### Immunohistochemistry

The cryosections were blocked with 1% skim milk in PBS with 0.5% Tween-20 at RT for 15 min and incubated with primary antibody (rat anti-mouse anti-PECAM-1 [CD31; BD Pharmingen, 553370; 1:400] or mouse monoclonal anti-QH-1 [Developmental Studies Hybridoma Bank, AB_531829; 1:100]) at 4°C overnight. The sections were washed in PBS four times at RT. For observation under an optical microscope, we incubated the sections with biotin-conjugated goat anti-rat IgG (Vector Laboratories, BA-9400) for CD31, and with goat anti-mouse HRP (Dako, P0447) for QH-1. For the fluorescent images, we incubated the sections with goat anti-rat IgG (Alexa Fluor 488) (Abcam, ab150165; 1:400) for 1 h. The sections were washed in PBS five times and mounted with Aqua-Poly/Mount (Cosmo Bio Co., Ltd., Japan). Nuclei were visualized with DAPI (Molecular Probes). Fluorescent signals were visualized under a computer-assisted confocal microscope (Nikon D-Eclipse C1), and images were processed using Nikon NIS Elements software (Nikon). The same protocol was followed for whole-mounted hearts and embryos, except that incubation with antibodies was performed overnight. For the whole-mount staining of the quails, we used biotin-conjugated goat anti-mouse IgG (Vector Laboratories, BA-9200) as the second antibody for CD31. The biotin-conjugated secondary antibodies were visualized using the Vectastain ABC System (Vector Laboratories, Burlingame, CA, USA). In the mouse histological sections in Figure 3, the peripheral nerves were visualized by using antibodies anti-acetylated tubulin (Sigma-Aldrich, no. T7451) and HRP-conjugated polyclonal goat anti-mouse (Dako, no. P0447).

### Ink and latex injection

To visualize the coronary arteries, we anesthetized *X. laevis*, red-bellied newts, and American bullfrogs at appropriate stages on ice or with 0.1% tricaine methanesulfonate (MS-222; Wako, Japan). Ink (Kiwa-Guro, Sailor, Japan) was gently injected using a glass micropipette from the hepatic portal vein. Then we collected their hearts, and the samples were fixed in 4% paraformaldehyde overnight at 4°C. The samples were cleared by CUBIC solution (Susaki et al., 2014) and examined under a stereo microscope. Mouse and quail hearts were injected with latex or resin as in Higashiyama et al., (2016b).

### Three-dimensional reconstructions of zebrafish samples

The confocal microscope images (z-stack: 2–5 μm) were used. For this images, the samples were fixed in 4% PFA overnight at 4°C, trimmed and immersed in CUBIC solution overnight at RT. Fluorescent signals were visualized under a Nikon D-Eclipse C1 microscope. The images were processed using Nikon NIS Elements software and loaded into the Amira 3D Visualization Software (Thermo Fisher Scientific) with a voxel size appropriate to section thickness.

### Live imaging of zebrafish

A multi-photon excitation microscope (FV1000MPE, Olympus) equipped with a water immersion 20× lens (XLUMPlanFL, 1.0NA, Olympus) was used. Sequential images were processed with FV10-ASW 3.1 viewer (Olympus) and analyzed with the Imaris software (Bitplane).

## DATA AVAILABILITY

All data generated or analysed during this study are included in the manuscript and supplementary information. The genomic data of *flk1:egfp* zebrafish is available in ZFIN; Tg(*kdrl:egfp*)^s843^, ID: ZDB-ALT-050916-14 (https://zfin.org/action/feature/view/ZDB-ALT-050916-14).

## ACKNOWLEDGEMENTS

We greatly appreciate Mayuko Kida and Akiyasu Iwase (The University of Tokyo) for discussion. We are also grateful to Japanese fish markets, Yoshiike (Tokyo, Japan), Induka Shouten (Nagasaki, Japan), and Gyosai-ya (Tokyo, Japan), for providing the chondrichthyan samples. We also appreciate Yuriko Kondo (The University of Tokyo) for secretarial assistance. This work has supported by the Grant-in-Aid for the Japan Society for the Promotion of Science (JSPS) 20H04858 and 20K15858 (H.H.).

## Author contributions

K.M., H.H., and H.K. conceived the study and designed experiments. K.M., H.H., Y.A., and S.F. performed experiments. S.F., KA. and S.M.T. supported the experiments. K.M., H.H., Y.A., and H.K. analyzed the data. K.M., H.H., and H.K. wrote the manuscript with help from other authors.

## SUPPLEMENTARY MATERIALS

**Fig. S1:**
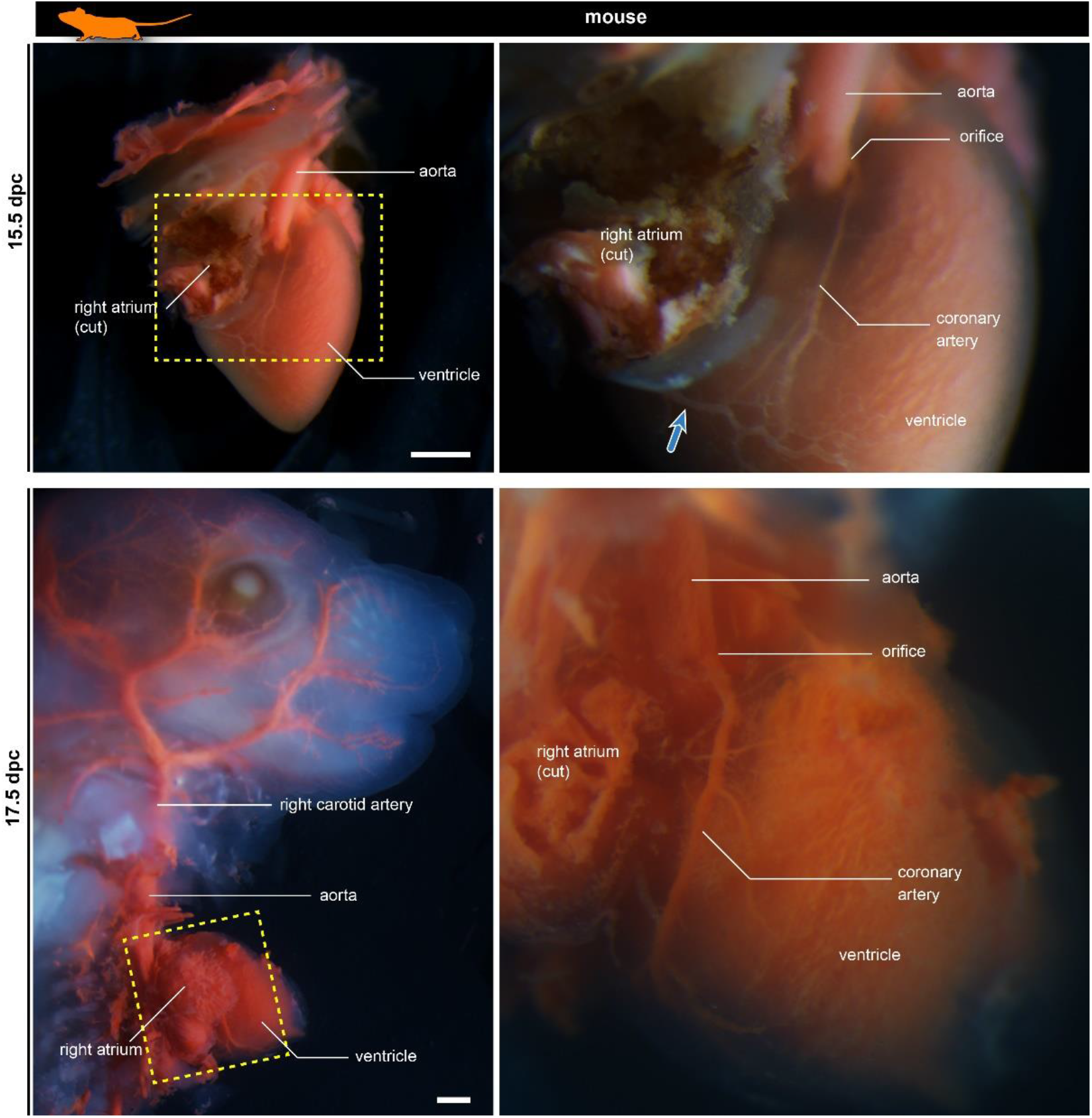
Development of murine coronary artery visualized in ink-injected fetuses. The blue arrow indicates the coronary vein. The right panels are the partial enlargements of the left panels. The blue arrow shows the coronary vein. Scale bars: 500 μm.

**Fig. S2:**
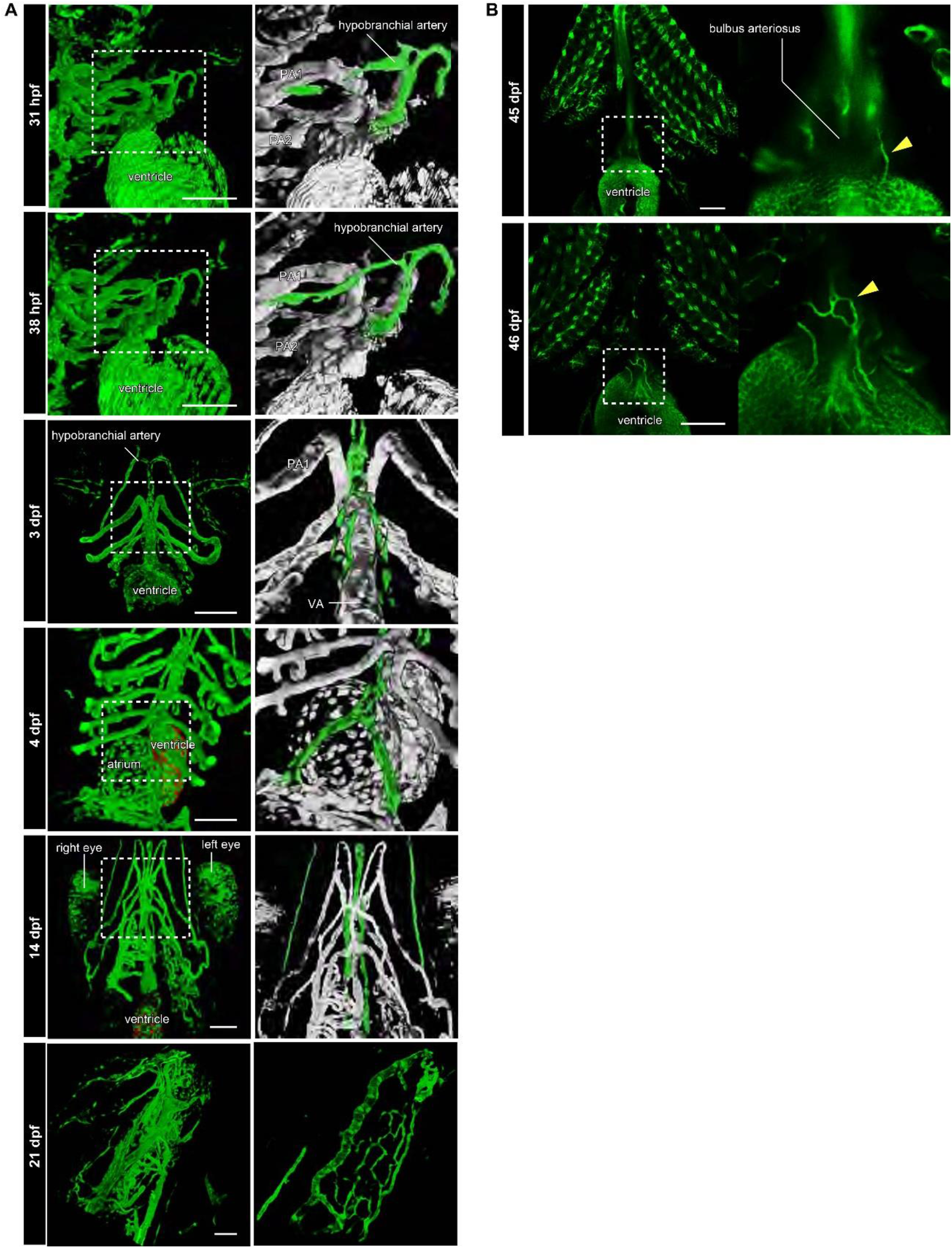
The development of hypobranchial artery in *flk1:egfp* zebrafish. **(A)** Two-photon excitation microscopy images of the heart and aortic arch. The hypobranchial arteries are green in the magnified images (right column). At 3–4 dpf, an *egfp-positive* endothelial bud appeared in the midline of the pharyngeal region and formed a Y-shaped structure. After 3 dpf, the hypobranchial artery expanded toward the cardiac outflow tract (ventral aorta). At 21 dpf, the hypobranchial artery formed a vascular plexus along the ventral aorta. **(B)** Confocal images of the ventral side of the heart. At 45 dpf, *egfp-positive* vessels formed a network on the surface of the outflow tract (bulbus arteriosus). At 46 dpf, these vessels connected with the hypobranchial artery. PA1, first pharyngeal artery; PA2, second pharyngeal artery; VA, ventral aorta. Scale bars: 100 μm.

